# Intracellular signaling in proto-eukaryotes evolves to alleviate regulatory conflicts of endosymbiosis

**DOI:** 10.1101/2023.07.05.547817

**Authors:** Samuel H. A. von der Dunk, Paulien Hogeweg, Berend Snel

## Abstract

The complex eukaryotic cell resulted from a merger between simpler prokaryotic cells, yet the relative timing and the role of the mitochondrial endosymbiosis with respect to other eukaryotic innovations has remained under dispute. Although expansion of the regulatory repertoire has been inferred from phylogenetic studies, gene regulation has not been taken into account in current scenarios of the mitochondrial endosymbiosis which mostly focus on the complementary energetic and ecological perspectives. The endosymbiotic state introduced several unique challenges to cells such as coordination of host and symbiont cell cycles and its disruption by leaking gene products and DNA fragments between host and symbionts. To investigate how these unique challenges impacted genome and network evolution during eukaryogenesis, we study a constructive computational model where two simple cells are forced into an obligate endosymbiosis.

Across multiple *in silico* evolutionary replicates, we observe the emergence of different mechanisms for the coordination of host and symbiont cell cycles, stabilizing the endosymbiotic relationship. The most commonly evolved mechanism, implicit control, works without signaling between host and symbiont. Signaling only evolves under the influence of leaking gene products, while such regulatory interference is inherently harmful. In the fittest evolutionary replicate, the host controls the symbiont cell cycle entirely through signaling, mimicking the regulatory dominance of the nucleus over the mitochondrion that evolved during eukaryogenesis.

## Introduction

The mitochondrial endosymbiosis was an important step in eukaryogenesis, which gave eukaryotes a crucial energetic benefit and increased evolutionary potential (Sagan, 1967; Lane and Martin, 2010; Eme et al., 2017; López-García et al., 2017). Yet the impact of the mitochondrial endosymbiosis on eukaryogenesis reaches beyond increased metabolic capacity. Through endosymbiotic gene transfer, the large majority of mitochondrial genes ended up in the nuclear genome contributing to its expansion (Timmis et al., 2004; Gabaldón and Huynen, 2007). In addition, the nucleus evolved regulatory control over mitochondria, subjugating them to the eukaryotic cell cycle (e.g. Scarpulla, 2011). Several other eukaryotic innovations—including the nucleus, intron splicing and sex—have also been posited to be direct consequences or adaptations to the mitochondrial endosymbiosis (Koonin, 2006; Lane, 2006, 2014; Raval et al., 2022), although the mitochondrial impetus for the evolution of the nucleus and introns has recently been called into question (Vosseberg et al., 2023). The mitochondrial endosymbiosis was unique relative to virtually all known subsequent endosymbioses (including the chloroplast) as these involved a eukaryotic host which had already adapted to endosymbionts (i.e. mitochondria). These mitochondria-carrying eukaryotic hosts feature a well-defined subcellular organization as well as several distinct regulation and signaling mechanisms to coordinate endosymbiosis. Likely as a consequence, eukaryotes commonly engage in endosymbiosis of prokaryotes and of other eukaryotes (Yoon et al., 2006; Keeling, 2013; Hehenberger et al., 2019). Prokaryotes, on the other hand do not feature subcellular organization and frequently engage in *ecto*symbiotic but not in *endo*symbiotic relations (with one notable exception; Von Dohlen et al., 2001). An endosymbiosis between two prokaryote-like organisms presents many challenges, such as cell-cycle coordination of endosymbionts, and preservation of genomic and functional integrity of the host despite leakage of symbiont molecules. Identifying mechanisms by which evolution can solve these challenges is important for understanding eukaryotic life. Modeling is uniquely suited to obtain insights into how evolution could potentially have overcome the challenges of endosymbiosis. As outlined, eukaryogenesis and endosymbiosis are complicated processes that involve multiple levels of organization, e.g. genomic, regulatory, cellular, holobiont. On top of that, eukaryogenesis occurred only once, roughly 1–2 billion years ago, leaving little direct data to probe the evolutionary forces that were at play (Betts et al., 2018). We recently developed a multilevel model based on cell-cycle regulation, which is well suited to investigate endosymbiosis in an evolutionary context (Von der Dunk et al., 2022). The cell cycle is a fundamental task of a cell which connects many levels of organization: from specific regulatory products and binding sites on the genome to the overall behavior of the cell (growth, replication, division). Furthermore, the cell cycle is likely a focal point of evolutionary changes in an endosymbiotic context, where host and symbiont need to coordinate their growth relative to each other. From a regulatory perspective, symbiont coordination is similar to cell size control, for which cells need to coordinate division with volume growth. Multiple strategies for cell size control are observed in nature, including sizers which divide when reaching a fixed volume, and timers which divide after a fixed amount of time (Sauls et al., 2016; Witz et al., 2019; Proulx-Giraldeau et al., 2022). Analogously, we here investigate how symbiont control is achieved. Using our multilevel model, we investigate how an endosymbiosis event like the mitochondrial endosymbiosis can impact cells at the level of genome, regulatory network, and cell-cycle dynamics.

## Methods

### Cell-cycle regulation in host and symbiont

To model obligate endosymbiosis including molecular interference between host and symbiont, we use as a basis our previous model of obligate endosymbiosis and cell-cycle regulation (von der Dunk et al. (2022), which is an extension of Von der Dunk et al. (2022) and Quiñones-Valles et al. (2014); see Fig. 1). Hosts and symbionts are modeled as entities that regulate an autonomous cell cycle consisting of four states including an S-phase, in which the genome has to be replicated, and an M-stage, in which cells divide. The timely expression of cell-cycle stages is achieved by a gene regulatory network that is formed by the interactions between regulatory genes (coding for regulatory products such as transcription factors) and binding sites encoded on a linear genome. Interactions are stochastic, with at most one product binding per binding site each time step. The binding probabilities are defined by bit string similarity using thermodynamic approximations (see von der Dunk et al., 2022). In the case of regulatory interference (i.e. product leakage between host and symbiont), foreign products enter the gene regulatory network and compete with native products for binding sites on the genome. The gene regulatory network evolves through mutations in sequences and other properties of genes and binding sites (regulatory effect, activation threshold), as well as through duplications, deletions, innovations and relocations of genes and binding sites on the genome.

**Figure 1:**
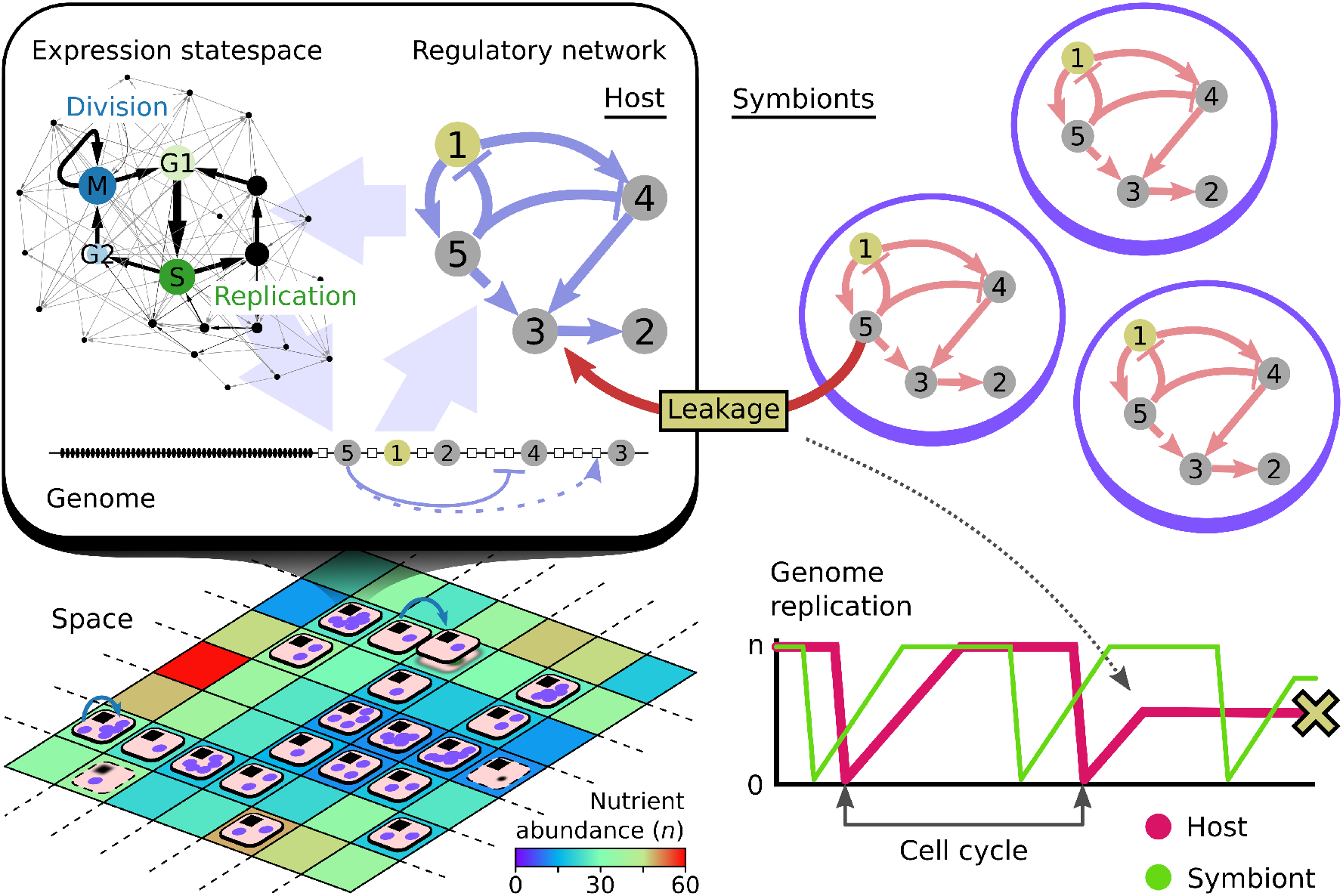
Overview of the modelling framework (cf. von der Dunk et al., 2022). Holobionts which consist of a host and one or more symbionts live on a grid where they compete for nutrients (bottom left). At the core of each host and symbiont is a genome with beads representing regulatory genes, binding sites and passive household genes (top left). Interactions between gene products and binding sites give rise to the regulatory network and expression dynamics constituting the cell cycle. The host and symbionts interact through product leakage, product targeting (i.e. signaling) and gene transfer. As an example, leakage of symbiont g5 product to the host is shown and the impact on cell-cycle dynamics: the host gets stuck mid-way through replication and later dies trying to divide with a partially replicated genome. In this and subsequent figures, regulatory interactions mediated by products that originate from the host are colored blue and those that originate from the symbiont are colored red.

### Balancing symbiont number with nutrient availability

Obligate endosymbiosis is modeled as holobionts consisting of one host and one or more symbionts, which live on a two-dimensional grid with an externally provided nutrient gradient (Fig. 1). The holobiont dies when the host dies or when there are no symbionts left, and the holobiont replicates into a new grid site when the host divides. Symbionts are stochastically distributed over offspring, so holobionts need to maintain more than one symbiont at the time of division to ensure that each daughter receives at least one symbiont. At the same time, hosts and symbionts share nutrients in the environment (3-by-3 neighborhood), so higher symbiont number leads to lower nutrient availability. Nutrients are required for replication, defining how many genomic elements (regulatory genes, binding sites, and passive household genes) are replicated during one time step in S-phase. Thus, to achieve fast growth, holobionts should not maintain too many symbionts.

### Product leakage and gene transfer before host compartmentalization

To study the evolutionary challenges of endosymbiosis, we model interference between the gene regulatory networks and genomes of host and symbiont, representing the holobiont before the emergence of structures or mechanisms that act as boundaries, such as the nucleus (Fig. 1). Molecular interference is implemented in both directions as to remain agnostic to the specific mechanisms of interference, although symbiont death would make symbiont-to-host interference more likely than the reverse.

The products of expressed genes leak between symbionts and the host with a rate of *l* = 0.01 per gene per symbiont. As a consequence, the influx into each symbiont is proportional to the number of expressed genes in the host, whereas the influx into the host is proportional to the number of expressed genes in all symbionts. Foreign products act like native products in all respects: they can bind to binding sites and they also define the cell-cycle stage if they are identical to one of the native core gene types (g1–5). Thus, product leakage disrupts expression dynamics and hinders the initially autonomous cell-cycle regulation of hosts and symbionts (Fig. 1).

Besides passive molecular interference, we include mutations that generate or destroy signal peptides of products (*μ*_*s*_ = 0.00001) yielding active targeting to the host or symbiont. The signal peptide for host and symbiont localization are represented by two bits, respectively: 10 defines host localization, 01 symbiont localization, 00 no relocation, and 11 dual localization. In the initial holobiont, all gene products are only targeted to the genome where they are encoded (10 or 01).

At the genome-level, interference consists in gene transfer between host and symbiont, which can disrupt the encoded gene regulatory network or foster innovation. A newly divided host (symbiont) can receive gene transfers from all symbionts (from the host) at a per-gene rate of *μ*_*T*_ = 0.00001, and we included both copy-and-paste and cut-and-paste type transfers.

## Results

### Holobionts adapt despite molecular interference

To study host and symbiont evolution in obligate endosymbiosis, we used our multilevel model of the cell cycle in the context of gene regulation (see Methods). In short, genomes undergo DNA replication which has to be timed with cell division. Hosts and symbionts have to evolve their cell cycle to adapt to poor and fluctuating nutrient conditions which requires prolonged duration of the S-phase. At the same time, the holobiont must also evolve coordination of symbiont division to avoid losing the obligate symbiont.

We evolved 10 replicate populations (Q1–10) with these endosymbiotic challenges, i.e. product leakage, gene transfer and signal peptide mutations. All replicates start with the same simple host and symbiont genomes, derived from Quiñones-Valles et al. (2014) (see also Von der Dunk et al. (2022); von der Dunk et al. (2022)). Despite initial identical genomes and gene regulatory networks, host and symbiont cell cycles do not stay synchronized due to stochasticity in the regulatory dynamics.

Adapting to the endosymbiotic lifestyle with these challenges is difficult, as we describe in detail for the most successful replicate as measured by population size, Q4 (Fig. 2; see Fig. S3–4 for two alternative trajectories). Early in the experiment, the population is small and confined to the richest environments on the gradient. A small population is prone to extinction, as seen in 3 other replicates (Q1, Q2 and Q6; Fig. 3). Moreover, selective forces are weak, so genome sizes drift extensively: in the first 10^6^ timesteps of Q4, the symbiont genome expands to 186% of its initial size (i.e. from *L* = 64 to *L* = 119.3). It takes until around *t* = 10^6^ for holobionts to become better adapted, resulting in rapid expansion of the population into poorer nutrient conditions on the gradient. At this time point, host and symbiont genome sizes no longer drift and decrease slightly as there is now strong selection for shorter genomes that need less time for replication and allow execution of a faster cell cycle. Subsequently and concomitant with additional population growth, both host and symbiont genomes expand again between *t* = 3 · 10^6^ and *t* = 5 · 10^6^ and reach a similar size. Surprisingly, the symmetry in genome size is retained for the rest of the experiment, with both genomes simultaneously shrinking or expanding at various occasions. Still, inspection of host and symbiont genomes reveals that they evolved encoding of different functions: the host has a large regulatory repertoire (*R*) whereas the whereas the symbiont is highly reduced in terms of regulation (Fig. 2). Vice versa, the symbiont in Q4 encodes most of the passive household genes (*L* — *R*) for the holobiont, indicating that there is specialization of regulation and other (non-regulatory) functions between host and symbiont.

**Figure 2:**
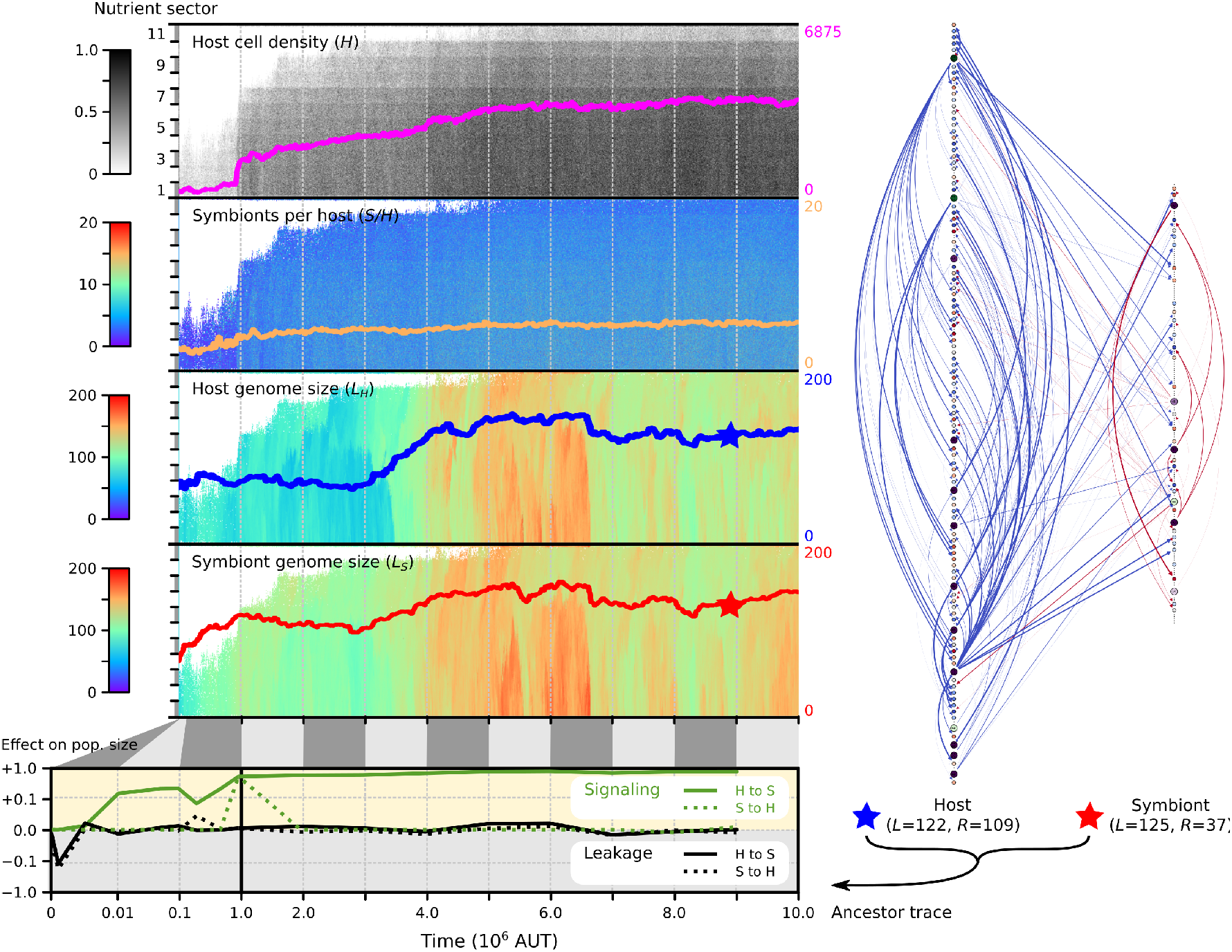
Evolution of host–symbiont signaling in holobionts adapting to a nutrient gradient (replicate Q4; see Fig. S3–4 for the evolutionary dynamics in two other replicates). The four panels at the top show the evolution of population size, symbiont number and genome size in space (the gradient with 11 nutrient sectors shown in the vertical dimension) and time (in the horizontal dimension). Overlaid on each space-time plot are the population averages across space. On the right, the genome networks are shown of the last common ancestors of all hosts and symbionts in the final population (marked with a star). The symbiont genome appears smaller than the host genome because it encodes mostly passive household genes (which are represented by small squares). In the bottom panel, we assessed the impact of leakage and signaling along the ancestral lineage. The first 10^6^ AUT are shown on a semi-log scale, revealing very rapid evolution of signaling and insensitivity to leakage.

**Figure 3:**
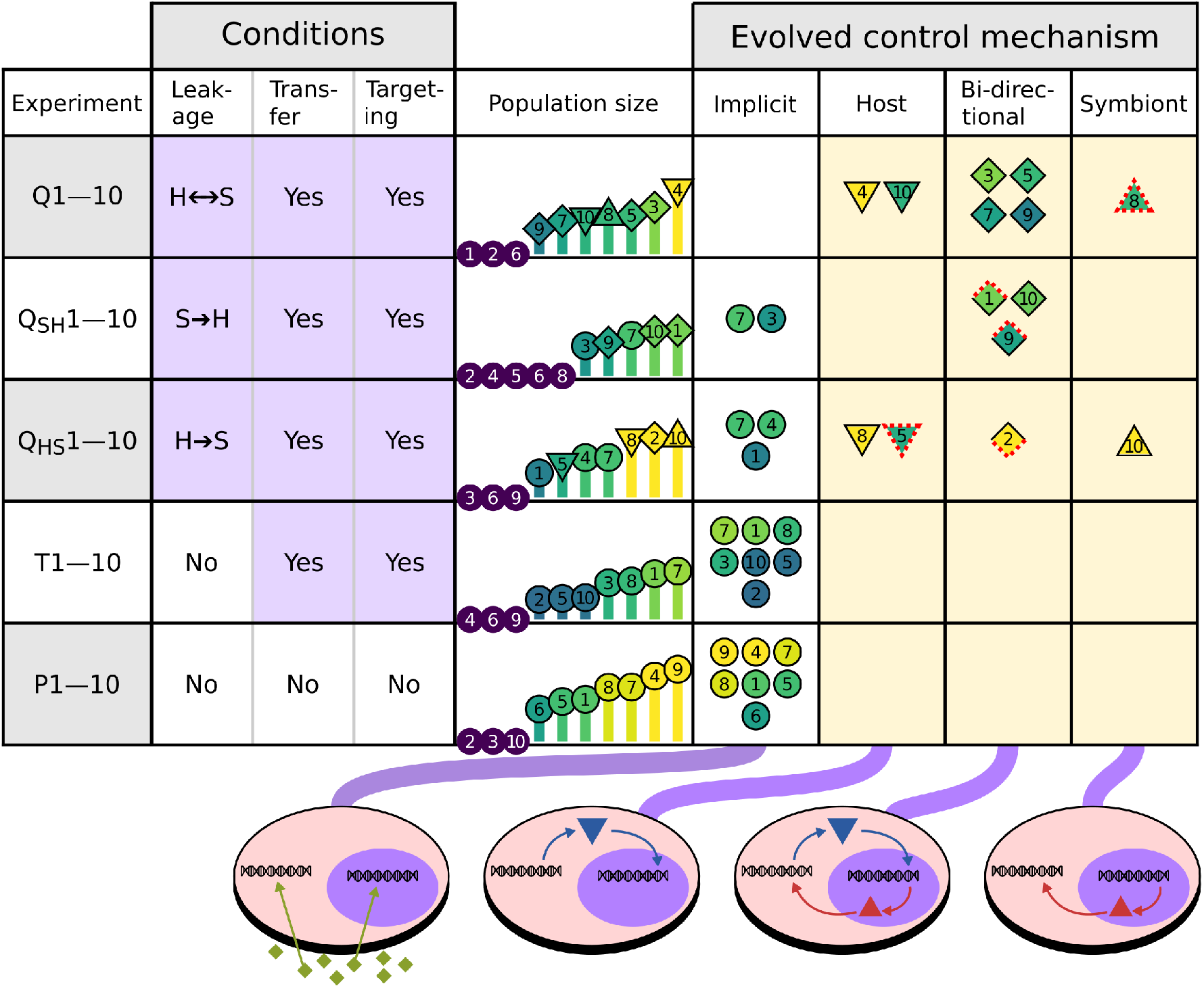
Leakage drives evolution of signaling, as shown by outcomes of five evolution experiments performed with different leakage and transfer conditions. With bi-directional leakage, signaling always evolves (creme shading; see cartoons on the bottom). With uni-directional leakage, signaling evolves in some replicates. Without leakage, only implicit control evolves, i.e. cell-cycle coordination without regulatory communication. Triangles indicate the direction of control (down for host control, up for symbiont control, diamond for bi-directional control), and red dotted lines indicate communication through leakage (down for host-to-symbiont direction, up for symbiont-to-host direction).

### Evolution of host–symbiont communication

As we set out to study the effects of molecular interference during endosymbiosis, we next investigated how holobionts adapted to product leakage along the ancestral lineage of Q4 (Fig. 2). To this end, clonal growth experiments were performed under high nutrient conditions (*n*_*influx*_ = 90) with bi-directional, uni-directional or no product leakage between host and symbiont. For the first ancestors, carrying capacity of clones grown in the presence of leakage is reduced compared to clones grown in the absence of leakage, confirming that product leakage is deleterious for early holobionts. Yet Q4 holobionts have rapidly evolved to be largely insensitive to leakage: their carrying capacity is the same in presence as in the absence of leakage (i.e. by *t* = 0.005 · 10^6^, bottom panel in Fig. 2).

In parallel with the leakage assay, we also investigated whether host–symbiont signaling evolved along the ancestral lineage of Q4. During the experiment, genes occasionally obtain signal peptides through mutations such that the gene product would be targeted from host to symbiont or from symbiont to host. However, many of the genes where this happens are not expressed and their cross-genomic interactions are not conserved through evolution. To detect whether *functional* signaling evolved, we studied how holobiont growth changes when targeting in one or both directions is blocked. Surprisingly, host-to-symbiont signaling emerges very early and has already become essential for the holobiont by *t* = 0.05 · 10^6^ (Fig. 2). Functional symbiont-to-host signaling appears transiently at *t* = 10^6^ but otherwise plays no role in holobiont growth. Thus, in Q4, passive communication in the form of product leakage is rapidly supplanted by active communication in the form of host–symbiont signaling. Signaling by the host establishes regulatory control over all six symbiont regulatory genes, allowing the host to order symbiont division and effectively manage symbiont number. We refer to this particular evolved strategy as host control.

### Product leakage drives evolution of signaling

Our modeling approach enables parallel evolutionary runs, which helps us to evaluate the generality of the evolution of host control as a mechanism to stabilize endosymbiosis. The 7 evolution replicates that survived and adapted to the nutrient gradient all evolved some form of signaling between host and symbiont (Fig. 3). Besides host control as in Q4 (and also in Q10), we found symbiont control mediated by symbiont-to-host signaling (Q8), and bi-directional control mediated by two-way signaling (Q3, Q5, Q7 and Q9; see Fig. S5–9). To study whether product leakage or gene transfer played a role in the invention of signaling, we performed several additional evolution experiments: with uni-directional leakage (Q_HS_1–10 and Q_SH_1–10), and without leakage altogether (T1–10). We also included in this comparison an earlier experiment without any interference or possibility for communication, i.e. no leakage, no transfer and no signal peptide mutations (P1–10; featured in von der Dunk et al. (2022)). Strikingly, signaling does not evolve in the absence of product leakage, despite mutations that generate signal peptides (T1–10). Moreover, under uni-directional leakage (Q_HS_1–10 and Q_SH_1–10), signaling evolves in some but not all replicates. These observations firmly establish that product leakage is what drives the evolution of host–symbiont signaling in our model. To understand why the evolution of signaling is correlated with product leakage, we must first examine the different evolved control mechanisms in more detail.

### Competitive success of evolved holobionts

With signaling, Q4 holobionts achieve high stability and large population size. Yet, holobionts in P9 and P4 also reach large population size with a strategy to balance host and symbiont growth, despite the fact that no signaling could evolve in these replicates. Population size reflects the fitness of individual holobionts (*R*_0_), but competitive success also depends on their generation time (i.e. at what timescale *R*_0_ is realized) and interaction with the environment (i.e. growth depends on nutrient levels which in turn depend on population size and symbiont numbers). To compare evolved holobionts and their strategies directly, we performed various competition experiments between populations. It turns out that holobionts from Q4 outcompete all other holobionts, because Q4 holobionts perform both accurate (high *R*_0_) and fast cell cycles. Nevertheless, other holobiont populations that evolved signaling (Q3,5,7–10 and Q_HS_2,10) are outcompeted by P4 and P9, showing that signaling is not required for competitive success.

One other important factor for competitive success of holobionts is the efficiency of individual host and symbiont cell cycles. At the start of the evolution experiment, host and symbiont regulate a primitive cell cycle. It takes substantial rewiring of the gene regulatory network to evolve a long and efficient cell cycle that can cope with poorer environments on the nutrient gradient (Von der Dunk et al., 2022). In different replicates, hosts and symbionts achieve different levels of adaptive rewiring, and this translates to different levels of success in holobionts. For instance, host and symbiont cell cycles are more efficient in P9 than in Q3 (i.e. higher *e*_*H*_ and *e*_*S*_ in P9; see panels b and d in Fig. 4), and so P9 is more successful in competition.

**Figure 4:**
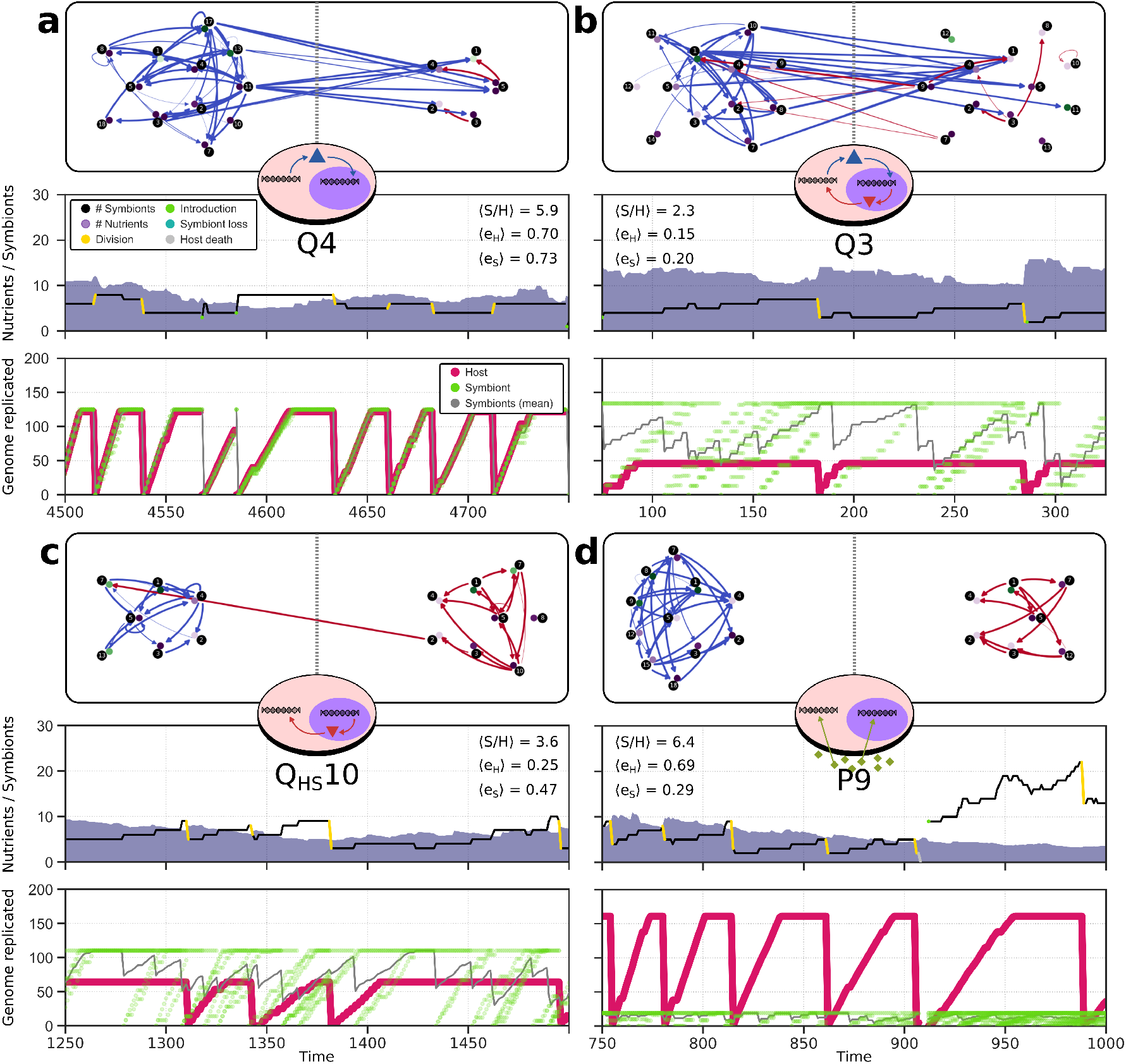
Four unique evolved mechanisms of cell-cycle coordination: (a) host control, (b) bi-directional control, (c) symbiont control, and (d) implicit control. For each mechanism, the most recent common ancestor of a successful replicate is analyzed in detail. The gene regulatory network of the holobiont is shown at the top, with regulation by host genes in blue and regulation by symbiont genes in red. Cell-cycle behaviour is analyzed in clonal growth experiments under intermediate nutrient conditions (*n*_*influx*_ = 30). In these growth experiments, the average symbiont number (*S/H*) and regulatory efficiencies of host and symbiont (*e*_*H*_ and *e*_*S*_) were also measured (see von der Dunk et al., 2022). High cell-cycle efficiency of the host, as seen in smooth replication trajectories (as opposed to more step-like), is partly responsible for the high fitness of Q4 and P9. The genome sizes of host and symbiont are clearly visible as the maxima in the replication panels.

### Mechanisms of cell-cycle coordination

Our modeling approach provides us with the opportunity to investigate alternative outcomes of endosymbiosis and therewith contextualize the single outcome of eukaryogenesis that appeared 1–2 billion years ago. Furthermore, our modeling approach allows us to pinpoint how different signaling mechanisms and control strategies work. To this end, we shall describe in the following sections for each holobiont strategy one successful replicate in terms of genome and network organization, regulatory behavior and cell-cycle dynamics (Fig. 4). Cell-cycle dynamics were distilled during a clonal growth experiment at intermediate nutrient condition (*n*_*influx*_ = 30) without product leakage. We followed a holobiont in a specific site on the grid, studying changes in symbiont number and in cell-cycle regulation of symbionts and the host through several holobiont divisions. Interestingly, the identified symbiont control strategies bear resemblance to different behaviors observed in cell size control (e.g. sizers and timers; see Supplementary Results).

### Host control achieves cell-cycle synchronization

As defined above, holobionts in the most successful replicate out of all experiments (Q4) evolved host control, resembling the outcome of eukaryogenesis and known secondary endosymbioses. Early on, several host products (starting with products of core genes g1 and g4) evolve dual localization, promoting the same expression dynamics in the symbiont as in the host. Over time, the symbiont copies of g1 and g4 lose their regulatory functions and these are taken over by the versions imported from the host. At the end of the experiment, only three important regulatory interactions are still carried out by the symbiont itself (Fig. 4a). The remaining symbiont genes (g1–5) are still functional as they define the symbiont’s cell-cycle stage. Moreover, these symbiont core genes have diverged from the core genes of the host, which prevents the latter from taking over and defining cell-cycle expression for the symbiont.

Host control results in synchronization of host and symbiont cell cycles (Fig. 4a), something that is often considered an early and important step in the evolution of endosymbiosis (Uchiumi et al., 2019). Synchronization ensures that each symbiont divides once for every host division, resulting in constant symbiont number during the life span of a holobiont. Because each symbiont divides once, new holobionts on average carry the same number of symbionts as their parents. Analogous to the timer strategy in cell size control, stochastic differences in symbiont number are inherited by the next generation of holobionts and symbiont number is strongly correlated between consecutive divisions (Fig. S2a). Stochastic differences in symbiont number, which arise from random distribution of symbionts at holobiont birth, are entirely corrected at the holobiont level. Holobionts with too few symbionts do not divide successfully and holobionts with too many symbionts grow slow due to low nutrient levels; both are outcompeted by holobionts that happened to receive an intermediate number of symbionts at birth.

Host control creates selection for equal genome sizes (as observed in Fig. 2). The holobiont life cycle is most efficient when replication of host and symbiont genomes takes the same amount of time, since division needs to be delayed by the host’s regulatory network until both genomes are fully replicated. Thus, cell-cycle synchronization promotes symmetry in genome size despite functional differentiation as observed before (Fig. 2).

### Bi-directional control tunes symbiont number

The most common signaling mechanism in our evolution experiment is bi-directional control (8 replicates; Fig. 3). The initial evolution of bi-directional control is similar to that of host control: host products obtain dual localization early in the experiment and start exerting regulatory control over symbiont gene expression. Subsequently, a symbiont product with host localization appears and fixes in the population. For example in Q3, the gene g9 is encoded on the symbiont and its product targeted to the host where it triggers cell-cycle progression towards M-stage, i.e. promoting division. In turn, g9 expression relies on activation from the symbiont-targeted host product g10 which binds the regulatory region upstream of g9 with low binding affinity. Interactions with low binding affinity are sensitive to gene dosage, meaning that replication of the host genome (increasing copy number of g10) and replication of symbionts (increasing copy number of g9) both increase the probability of host division. Moreover, g10 is located near the terminus of the host genome. Thus in Q3, the host has evolved a checkpoint that stalls the cell cycle until the host has finished genome replication and until there are enough symbionts to safely divide the holobiont.

Bi-directional control has exposed a different mechanism for control of symbiont numbers relative to synchronization under host control (Fig. 4b). Holobionts with bi-directional control display behavior that resembles aging: during the lifespan of a holobiont, symbiont number gradually increases resulting in further depletion of nutrients and slowing down of replication. When a holobiont is old and carries many symbionts, it divides and gives birth to two young holobionts carrying on average half the number of symbionts. Importantly, deviations from this average are corrected in the next cell cycle through the aforementioned division checkpoint. Thus, symbiont numbers are not controlled at the holobiont population level as in the case of host control, but at the level of individual regulation, i.e. through signaling between host and symbiont. Bi-directional control thereby resembles the sizer mechanism for cell size control (Fig. S2b).

Interestingly, in holobionts with bi-directional control, the host usually drives the symbiont cell-cycle to a large extent: in Q3, the host regulates 5 symbiont genes whereas the symbiont only regulates a single host gene. Thus, host and symbiont have both lost autonomy, but the host is more dominant in terms of regulation reminiscent of real endosymbiotic relationships.

### Symbiont control by two separate checkpoints for host division

The least common signaling mechanism that evolved is symbiont control (2 replicates; Fig. 3), and Q_HS_10 is the most successful replicate with this mechanism (Fig. 4c). In contrast to host-to-symbiont signaling (e.g. in Q3 and Q4), symbiont-to-host signaling arises very late in evolution (e.g. in Q3 and Q_HS_10). The symbiont gene product g2 evolves dual localization around *t* = 3 · 10^6^, but only establishes functionally relevant regulation of the host around *t* = 9 · 10^6^. Specifically, g2 targets and inhibits host g7, the gene product that stalls the host cell cycle in S-phase and delays division. Symbiont g2 is expressed infrequently during the symbiont cell cycle and rarely at the time that is required to promote host division. Higher symbiont number increases the probability that a g2 copy from any symbiont is expressed at the right time. Thus similar to bi-directional control, symbiont control acts to inform the host that there are enough symbionts to safely divide the holobiont, again analogous to a sizer in cell size control (Fig. S2c).

A key difference with bi-directional control in Q3 is that the interaction between symbiont g2 and host g7 in Q_HS_10 is not dosage-sensitive and does not measure the replication status of the host. The host has evolved a second mechanism to infer the replication status of its own genome. A gene near the origin, g13, performs the same regulatory tasks as g7, and thus prevents host division even if g7 is deactivated under the influence of high symbiont number. Instead, g13 needs to be inhibited by g5, which is located right at the terminus of the host genome. The g5–g13 interaction depends on low-affinity binding, so inhibition of g13 only becomes likely when the entire host genome is replicated (giving two copies of g5). Thus, where bi-directional control in Q3 integrates host status and symbiont levels into a single cell-cycle checkpoint, symbiont control in Q_HS_10 rests on two independent checkpoints for holobiont division.

### Product leakage exploited as additional channel for communication

Interestingly, we also found cases of host control, bi-directional control and symbiont control that rely on product leakage rather than signaling (see Fig. 3). For instance in Q8 (an outcome with symbiont control), hosts occasionally enter a dormant state in their cell cycle where no genes are expressed. In this state, they depend on the leakage of symbiont products to start up their cell cycle again. As the rate of product leakage scales with symbiont number, symbiont control through passive leakage and symbiont control through signaling both work by stalling the host cell cycle until there are enough symbionts to safely divide the holobiont.

In Q_HS_2 (a case of bi-directional control), the host leaks the product of gene g1 to the symbiont, where it activates gene g14 whose product is then actively targeted back to the host to induce cell-cycle completion. Thus, product leakage and signaling can operate in unison to achieve cell-cycle coordination. In the same way that low binding affinity sets a timescale on a regulatory interaction, the leakage rate also sets a timescale on regulatory events. Moreover, low binding affinity interactions and effective leakage rates are both sensitive to gene dosage (i.e. expression frequency, replication status, symbiont number) and can thus be used as cell-cycle checkpoints that integrate external information into the regulatory network. In this light, it is interesting to note that Q_HS_2 is much more successful than Q3 which uses signaling in both directions for bi-directional control.

We have seen that signaling does not evolve without leakage and that leakage itself can also be exploited for host–symbiont coordination. These observations could suggest that such functional leakage acts as an intermediate step in the evolution of signaling, whereby a host (symbiont) product first acquires a regulatory role in the symbiont (host) mediated by leakage and then later acquires the target signal. This is not the case: as we saw in Q4 (Fig. 2), leakage never performs a functional role in the holobiont prior to the emergence of signaling. It appears that the critical effect of leakage is that it compromises host and symbiont cell-cycle autonomy, which is vital for execution of the distinct host and symbiont cell cycles that accomplish implicit control (see below). Leakage thereby stimulates the integration of regulatory networks through communication to overcome its negative effect on implicit control, and this integration is achieved through active signaling or by exploiting leakage directly.

### Implicit cell-cycle control indirectly tunes symbiont number

The most common evolved coordination strategy—and the only outcome in the absence of product leakage—is implicit control, which we discovered as outcome in our previous study (von der Dunk et al., 2022). In this strategy, the host does not explicitly communicate with symbionts to decide when it divides. Instead, hosts and symbionts have evolved specialization on distinct nutrient conditions and form a stable equilibrium in their growth dynamics. Fluctuations in the local nutrient condition (indirectly reflecting changes in symbiont number) are countered by an increase or decrease in the growth rate of symbionts relative to the host until the system returns to the equilibrium state where host and symbiont grow at the same rate. It is remarkable that autonomous cell-cycle behavior of host and symbiont achieves such successful coordination, especially since nutrients only provide a coarse measure of symbiont number in the local environment. Moreover, in comparison to bi-directional control and symbiont control, implicit control takes several holobiont generations to correct high or low symbiont numbers resulting from asymmetric holobiont division. Because it takes multiple generations to correct deviations in symbiont numbers from the mean, implicit cell-cycle control behaves like a weak sizer (Fig. S2d).

In most replicates with implicit control, there is symmetry breaking in genome size between the host and the symbiont. This can be explained by selection on the relative cell-cycle speeds of host and symbiont. In P9 for instance, to achieve a stable holobiont with high symbiont number, the host evolves a slower cell cycle than the symbiont. The slow cell cycle of the host gives it more time to do replication and this is exploited by evolution through transfer of household genes from symbiont to host. We also found a bias in evolution towards a large host and small symbiont genome, resembling the outcome of eukaryogenesis, and it turns out that holobionts with such asymmetry perform best (von der Dunk et al., 2022).

### Historical contingency favors implicit control

All the experiments described so far involved identical initial host and symbiont genomes. In the light of eukaryogenesis, it is interesting to also consider the effects of molecular interference between a host and symbiont that are more distantly related, analogous to an Asgard archaeon and an Alpha-proteobacterium. For this, we exposed the 7 surviving replicate populations that evolved without leakage and transfer and signal peptide mutations (P1–10) to all of these processes from *t* = 10 · 10^6^ onwards (Fig. 5). The 7 replicate populations had all evolved implicit control to accomplish stable holobiont growth (Fig. 3). Three replicates evolved high symbiont number resulting in stable, slow-growing holobionts and four replicates evolved low symbiont number resulting in unstable, fast-growing holobionts.

**Figure 5:**
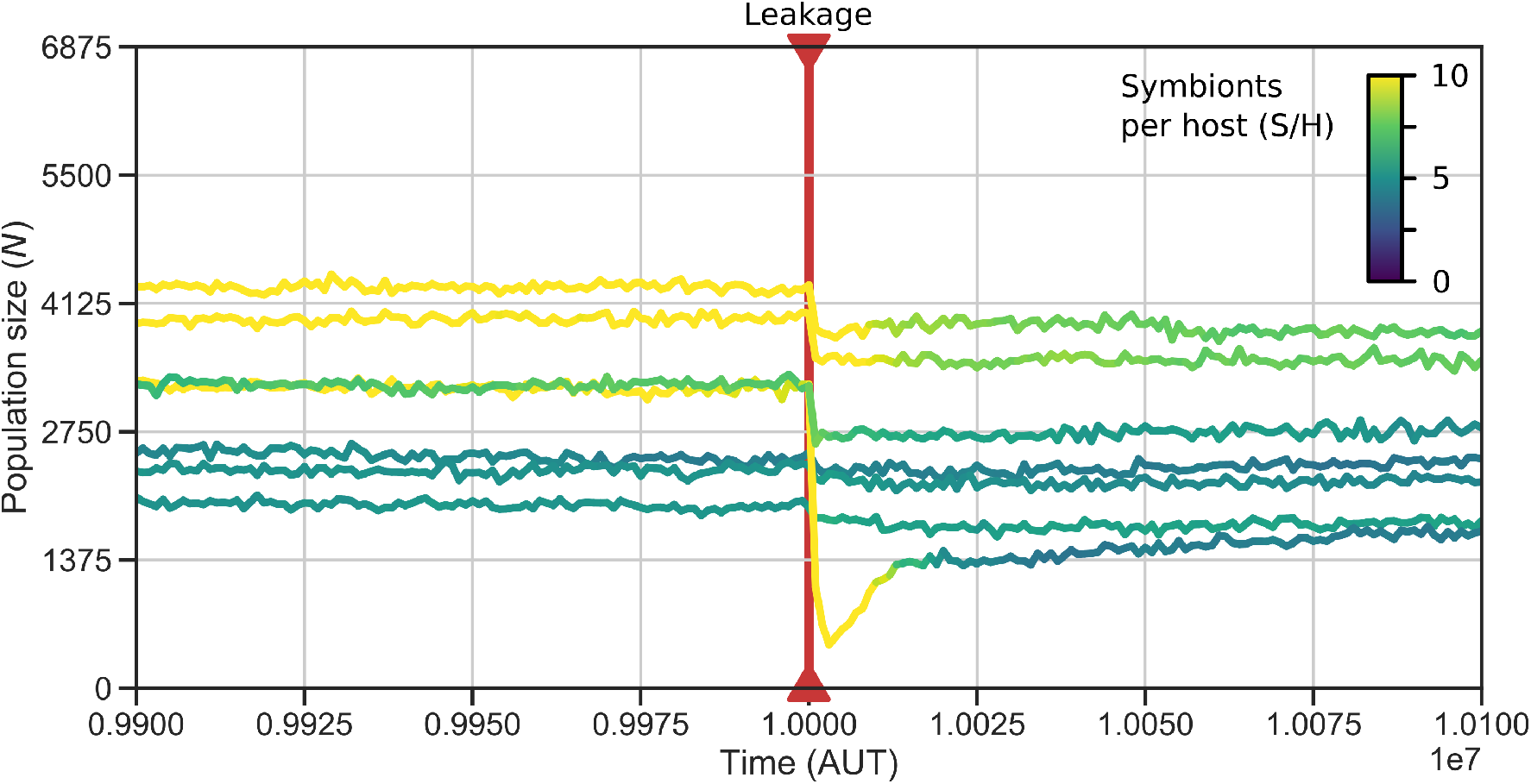
Product leakage hinders holobionts even after they are adapted to the endosymbiotic state and host and symbiont have diverged. The 7 surviving replicates of P1–10 were exposed to leakage, transfer and signal peptide mutations from *t* = 10 · 10^6^, causing immediate drops in population size. Populations which evolved high symbiont numbers are affected most: P7 nearly goes extinct. Holobionts readapt by decreasing symbiont number.

The introduction of molecular interference disrupts holobiont behavior resulting in immediate population size reductions (by up to 66.7% in the first 1000 timesteps). Thus, despite the genetic distance between host and symbiont, molecular interference still presents a hazard for correct regulation of their cell cycles. Product leakage has the biggest direct impact on holobiont growth, although gene transfer also impacts holobionts on a long timescale by altering genome size evolution (see Supplementary Results). Replicates that had evolved high symbiont number before the transition to the new regime are impacted more than those that had evolved low symbiont number, because effective leakage rates are higher in holobionts with more symbionts. Holobionts quickly adapt to product leakage by decreasing symbiont number. Yet, the mechanism of cell-cycle coordination does not change. The genetic distance between host and symbiont makes signaling difficult to evolve in our genotype-phenotype map. Implicit cell-cycle control does not require genetic integration and is therefore much easier to evolve between distantly related host and symbionts. The same outcome was also clearly visible in another set of experiments starting with pre-evolved free-living host and symbionts (see Supplementary Results). Thus, historical contingency, i.e. the evolutionary past of free-living cells before they engage in endosymbiosis, favors a fully autonomous host and symbiont which balance their nutrient-dependent growth rates.

## Discussion

### Evolution of signaling and regulatory dominance of the host

We investigated the impact of obligate endosymbiosis in simple cells that are not preadapted to the prevailing environment and in particular have never been exposed to the physical challenges of endosymbiosis. The main challenge of endosymbiosis is to control the symbiont population through cell-cycle coordination between host and symbiont. Additionally, product leakage disrupts the regulatory autonomy of host and symbiont. Here we show that product leakage is not only a challenge but in fact also drives the evolution of intracellular signaling, indirectly aiding the establishment of cell-cycle coordination and stable holobiont growth.

Out of the three different signaling mechanisms that arose in our evolution experiment, host control performed best in terms of population size and direct competition experiments. Host control has also been the outcome of eukaryogenesis and all known secondary endosymbiosis events. In the more recent endosymbiosis of *Paulinella chromatophora* and its chromatophores, we already see signs of host control, by loss of symbiont genes or transfer to the host genome (Gabr et al., 2022). Our model shows that alternative mechanisms and coordination strategies can arise, but that the mechanism found in nature corresponds to the most successful of all strategies. The reason for this is that host control, and the resulting host–symbiont cell-cycle synchronization, is very effective in controlling symbiont populations. It is surprising that host control which amounts to a timer in cell size control, is so effective: symbiont numbers are largely controlled by selection at the holobiont-level, constituting frequent holobiont death. In contrast, sizers, which appear in bi-directional control and symbiont control, accomplish control of symbiont numbers by individual holobionts and result in fewer holobiont deaths.

In all evolved control mechanisms, the host turns out to be dominant in terms of regulation. In replicates with bi-directional control, hosts generally regulate multiple symbiont genes whereas symbionts often regulate a single host gene. Among replicates with symbiont control, a single symbiont product is targeted to the host in one replicate (Q_HS_10), and only leakage from symbiont to host is exploited in the other replicate (Q8). These observations invoke the image of symbionts merely informing the host rather than completely controlling it. The host could potentially still re-adapt to a free-living lifestyle by removing its single dependency on regulation by the symbiont, whereas the symbiont has become entrenched in the host.

At the level of genome size, no general asymmetry between host and symbiont was found to evolve, unlike in our previous study (von der Dunk et al., 2022) and unlike eukaryogenesis. In our model, cell-cycle speed poses a major constraint on genome size, and to limit symbiont numbers and regulatory conflicts, symbionts evolve relatively slow cell cycles. However, genome size asymmetry has classically been explained from energetic (Lane and Martin, 2010) or mutational perspectives (Allen and Raven, 1996; Martin and Herrmann, 1998) rather than a purely informational one. These other perspectives could provide potential additional reasons for the evolution of genome size asymmetry between host and symbiont.

### Mechanistic insights into the evolution of signaling

Signaling is often assumed to be readily evolvable as an adaptation to obligate endosymbiosis. Yet in our model, signaling only evolves under stress from product leakage and with simple and identical initial host and symbiont genomes. The integration of gene regulatory networks becomes harder when host and symbiont genomes are very different or already execute efficient autonomous cell-cycles before being forced into an endosymbiotic relationship. In these cases, implicit control evolves rapidly because it only requires finetuning of the existing regulatory behavior to particular nutrient conditions. Interestingly, we had expected gene transfer to enable the evolution of signaling, but gene transfers are mostly deleterious and rarely fixed by selection. Signaling emerges by relocalization of existing products to the endosymbiotic partner. This indicates that is easier for a gene whose expression is already controlled by the cell cycle to evolve a new function (i.e. neo-functionalization of the gene product) than for a gene with a new function to evolve to be under proper cell-cycle control (i.e. neo-functionalization of the regulatory region). This is likely due to the fact that in our model, regulatory regions are relatively large (spanning several binding sites), such that adaptation of the regulatory region requires more mutations than adaptation of the gene itself.

Using a multilevel computational model and *in silico* evolution experiments, we have shown that cell-cycle coordination in holobionts can be achieved through implicit or through explicit control. Implicit control occurs at the metabolic level, i.e. through interaction of host and symbiont with nutrients in the external environment. Explicit control occurs at the gene regulatory network level, i.e. through signaling or leakage, which allows for more direct and tight control of symbiont numbers. In both cases, the regulatory repertoire of the host is generally dominant over the symbiont, even if it the host cell cycle is under regulatory control of the symbiont. The general picture that emerges is that an endosymbiotic relationship can be stabilized at multiple organizational levels, and even in case of harsh conflicts at the molecular level. From an informational viewpoint, stable coordination of endosymbiosis does not present a hurdle for simple cells (prokaryotes); rather, endosymbiotic challenges help to explain how regulatory complexity increased during eukaryogenesis, and in particular, the molecular conflicts between host and symbiont may explain the emergence of intracellular signaling in proto-eukaryotes.

## Acknowledgements

The authors gratefully acknowledge the help of Jan Kees van Amerongen for running the local computer cluster, and Bram van Dijk for proofreading the manuscript.

## References

Allen, J. F. and Raven, J. A. (1996). Free-radical-induced mutation vs redox regulation: costs and benefits of genes in organelles. Journal of Molecular Evolution, 42:482–492.

Betts, H. C., Puttick, M. N., Clark, J. W., Williams, T. A., Donoghue, P. C., and Pisani, D. (2018). Integrated genomic and fossil evidence illuminates life’s early evolution and eukaryote origin. Nature ecology & evolution, 2(10):1556–1562.

Eme, L., Spang, A., Lombard, J., Stairs, C. W., and Ettema, T. J. (2017). Archaea and the origin of eukaryotes. Nature Reviews Microbiology, 15(12):711–723.

Gabaldón, T. and Huynen, M. A. (2007). From endosymbiont to host-controlled organelle: the hijacking of mitochondrial protein synthesis and metabolism. PLoS computational biology, 3(11):e219.

Gabr, A., Stephens, T. G., and Bhattacharya, D. (2022). Loss of key endosymbiont genes may facilitate early host control of the chromatophore in paulinella. Iscience, 25(9):104974.

Hehenberger, E., Gast, R. J., and Keeling, P. J. (2019). A kleptoplastidic dinoflagellate and the tipping point between transient and fully integrated plastid endosymbiosis. Proceedings of the National Academy of Sciences, 116(36):17934–17942.

Keeling, P. J. (2013). The number, speed, and impact of plastid endosymbioses in eukaryotic evolution. Annual review of plant biology, 64:583–607.

Koonin, E. V. (2006). The origin of introns and their role in eukaryogenesis: a compromise solution to the introns-early versus introns-late debate? Biology direct, 1:1–23.

Lane, N. (2006). Power, sex, suicide: mitochondria and the meaning of life. Oxford University Press.

Lane, N. (2014). Bioenergetic constraints on the evolution of complex life. Cold Spring Harbor perspectives in biology, 6(5):a015982.

Lane, N. and Martin, W. (2010). The energetics of genome complexity. Nature, 467(7318):929–934.

López-García, P., Eme, L., and Moreira, D. (2017). Symbiosis in eukaryotic evolution. Journal of theoretical biology, 434:20–33.

Martin, W. and Herrmann, R. G. (1998). Gene transfer from organelles to the nucleus: how much, what happens, and why? Plant physiology, 118(1):9–17.

Proulx-Giraldeau, F., Skotheim, J. M., and François, P. (2022). Evolution of cell size control is canalized towards adders or sizers by cell cycle structure and selective pressures. Elife, 11:e79919.

Quiñones-Valles, C., Sánchez-Osorio, I., and Martínez-Antonio, A. (2014). Dynamical modeling of the cell cycle and cell fate emergence in caulobacter crescentus. PloS one, 9(11):e111116.

Raval, P. K., Garg, S. G., and Gould, S. B. (2022). Endosymbiotic selective pressure at the origin of eukaryotic cell biology. Elife, 11:e81033.

Sagan, L. (1967). On the origin of mitosing cells. Journal of theoretical biology, 14(3):225–IN6.

Sauls, J. T., Li, D., and Jun, S. (2016). Adder and a coarse-grained approach to cell size homeostasis in bacteria. Current opinion in cell biology, 38:38–44.

Scarpulla, R. C. (2011). Metabolic control of mitochondrial biogenesis through the pgc-1 family regulatory network. Biochimica et biophysica acta (BBA)-molecular cell research, 1813(7):1269–1278.

Timmis, J. N., Ayliffe, M. A., Huang, C. Y., and Martin, W. (2004). Endosymbiotic gene transfer: organelle genomes forge eukaryotic chromosomes. Nature reviews genetics, 5(2):123–135.

Uchiumi, Y., Ohtsuki, H., and Sasaki, A. (2019). Evolution of self-limited cell division of symbionts. Proceedings of the Royal Society B, 286(1895):20182238.

von der Dunk, S. H., Hogeweg, P., and Snel, B. (2022). Obligate endosymbiosis explains genome expansion during eukaryogenesis. bioRxiv.

Von der Dunk, S. H., Snel, B., and Hogeweg, P. (2022). Evolution of complex regulation for cell-cycle control. Genome biology and evolution, 14(5):evac056.

Von Dohlen, C. D., Kohler, S., Alsop, S. T., and McManus, W. R. (2001). Mealybug β-proteobacterial endosymbionts contain γ-proteobacterial symbionts. Nature, 412(6845):433–436.

Vosseberg, J., Stolker, D., von der Dunk, S. H., and Snel, B. (2023). Integrating phylogenetics with intron positions illuminates the origin of the complex spliceosome. Molecular Biology and Evolution, 40(1):msad011.

Witz, G., van Nimwegen, E., and Julou, T. (2019). Initiation of chromosome replication controls both division and replication cycles in e. coli through a double-adder mechanism. Elife, 8:e48063.

Yoon, H. S., Reyes-Prieto, A., Melkonian, M., and Bhattacharya, D. (2006). Minimal plastid genome evolution in the paulinella endosymbiont. Current Biology, 16(17):R670–R672.

