## Supplementary Material for "Intracellular signaling in proto-eukaryotes evolves to alleviate regulatory conflicts of endosymbiosis"

Samuel H. A. von der Dunk[1*], Paulien Hogeweg[1], Berend Snel[1]

[1]Theoretical Biology and Bioinformatics, Department of Biology, Science Faculty, Utrecht University

Padualaan 8, 3584 CH, Utrecht, The Netherlands

[*]

July 5, 2023

#### Supplementary Results

##### Initial host–symbiont differences promote implicit control

All the experiments described in the main text were initiated with identical host and symbiont genomes. In the light of eukaryogenesis, it is interesting to compare these results with evolution experiments starting with pre-evolved and diverged lineages of "prokaryotes". To this end, we setup an evolution experiment with 12 different host–symbiont pairs, picking different prokaryotes that already adapted to poor nutrient conditions in an earlier study (Von der Dunk et al., 2022) (see Supplementary Table S1–2). Each host–symbiont pair was evolved in two or three separate replicates to untangle the effect of specific pairings. To assess the impact of product leakage and gene transfer, we performed the whole experiment under different conditions: with leakage, transfer and signal peptide mutations (X1–12); with leakage only (L1–12); with transfer only (A1–12); and without leakage, transfer or signal peptide mutations (C1–12, see von der Dunk et al., 2022).

In the evolution experiments with different pairs of pre-evolved hosts and symbionts, product leakage and gene transfer have a negative impact on final population size (Fig. S1). Across the 32 replicates evolved under leakage, transfer and signal peptide mutations, holobionts either went extinct or evolved implicit control—rather than any form of

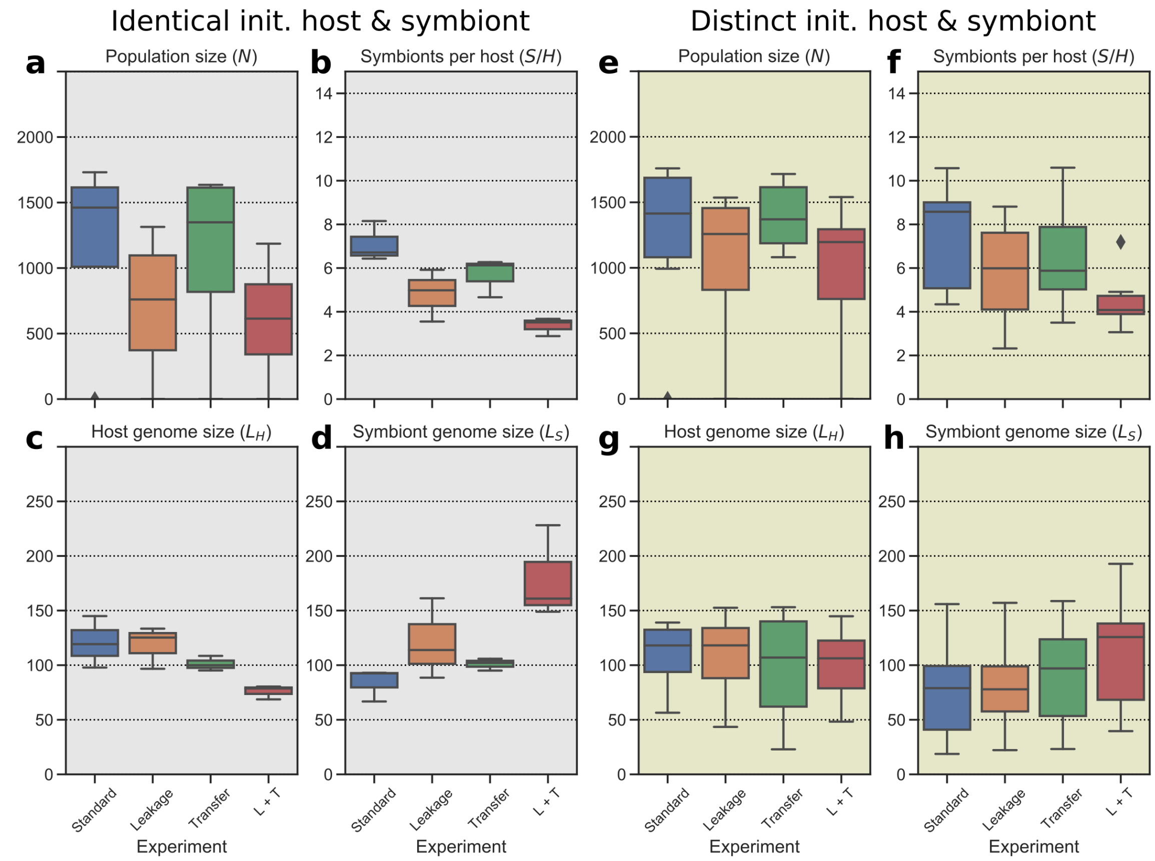


Figure S1: Product leakage and gene transfer are deleterious for holobionts consisting of pre-evolved host–symbiont pairs, as seen in smaller final population sizes (a,e) and larger genomes (c,g and d,h) at $t = 2 \cdot 10^6$. These effects can be explained by the reduced symbiont numbers (b,f) which holobionts evolved to limit the effective leakage and transfer rates from symbiont to host. The impact of leakage is most negative on holobionts initialized with identical host and symbiont genomes (a–d) compared to holobionts initialized with distinct host and symbiont genomes (e–h).

signaling—to coordinate host and symbiont cell-cycles. Thus, if we combine these results with those from the experiments described in the main text, implicit control is by far the most common evolved strategy in our model. Signaling only emerges when relatively primitive hosts and symbionts (i.e. which are not adapted to poor nutrient conditions) are exposed to product leakage from the start.

The inability to evolve signaling might be an important reason why holobiont populations remain smaller in the presence of product leakage in particular, even after having evolved for $t = 2 \cdot 10^6$ time steps. The main adaptation to product leakage, and to a lesser extent to gene transfer, is to limit symbiont number, reducing the effective rates of leakage and transfer. Symbiont numbers are limited by increasing the cell-cycle duration of the symbiont relative to the host, such that symbiont numbers are always diluted over multiple holobiont generations and are kept just above the bare minimum by holobiont-level selection. A slow symbiont cell cycle relative to that of the host relaxes selection for short symbiont genomes. Consequently, symbiont genomes expand more under the presence of leakage, and even grow bigger than host genomes. Interestingly, the small populations of symbionts with large genomes inside each holobiont are more sensitive to Muller's ratchet (von der Dunk et al., 2022). In particular, the formation and modification of regulatory genes on the symbiont genome is a hazard because these may subsequently leak or transfer to the host genome.

The impact of leakage and transfer is largest when evolution is started with identical host and symbiont genomes, i.e. similar to the situation in the experiments described in the main text. When host and symbiont genomes start out identical, genes likely retain some of their ancestral overlap, resulting in the strongest molecular interference. For example, the host copy of gene product g5 might still bind relatively strongly to binding sites for the symbiont copy of gene product g5. While many bit strings change substantially during evolution, we previously showed that bit strings of gene products that target more binding sites are more conserved than those of gene products with no or few regulatory targets (see "Gene family analysis" in Appendix 1 of Von der Dunk et al., 2022). Apparently, hosts and symbionts are unable to modify their gene regulatory network such that the effects of leakage are avoided at the level of gene regulation.

#### Symbiont control strategies

To understand how different control mechanism achieve stabilization of symbiont numbers, we tracked individual cells growing at intermediate nutrient conditions ($n_{influx} = 30$), either through their normal division process or by enforcing symmetric distribution of symbionts. By analogy to the literature on cell size control, wherein cell behavior can be analyzed by correlating the cell volumes between two consecutive cell cycles, we here compare the symbiont number right after division between two consecutive cell cycles (Fig. S2). Using this approach, the four different symbiont control strategies could be grouped into three different phenomenological control behaviors as mentioned in the main text. Host control ensures replication of all symbionts during the holobiont cell cycle, suggesting that deviations in symbiont numbers are propagated to subsequent generations. In line with this description, there is a high correlation between symbiont numbers of consecutive generations (Fig. S2a). Furthermore, holobiont deaths are much more frequent with host control than in any of the other control mechanisms, indicating that holobiont-level selection is important for stabilizing symbiont numbers in the population.

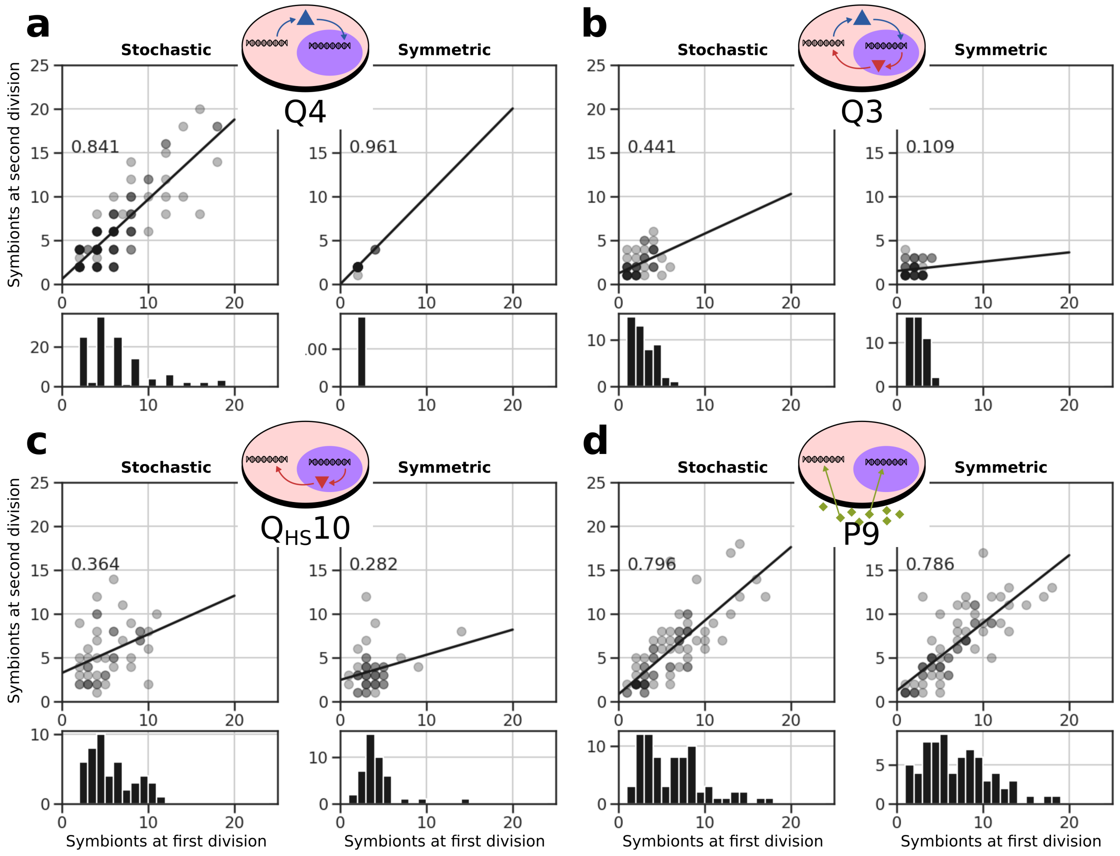


Figure S2: Phenomenological control strategies identified by correlations in symbiont numbers between consecutive cell cycles. As in Fig. 4 in the main text, the most recent common ancestor of each replicate was analyzed at intermediate nutrient conditions ($n_{influx} = 30$). Removing the stochasticity at cell division by forcing symmetric division shows the control strategy more clearly (right versus left panels). In the case of P9, the correlation stays well below 1, indicating that holobionts approach the equilibrium symbiont number over multiple generations. See also Fig. S10.

In contrast, bi-directional control and symbiont control establish division checkpoints that stall the cell cycle until sufficiently many symbionts are present. In line with this mechanism, the correlation between symbiont numbers in consecutive generations is small (Fig. S2b,c). Strikingly, the removal of stochasticity at cell division, i.e. by enforcing symmetric symbiont distribution, makes the control behaviors of host control on one hand, and of bi-directional control and symbiont control on the other hand, even clearer.

Finally, implicit control constitutes a weak sizer that takes several generations to correct deviations in symbiont number. For this reason, we do not find such a clear sizer signal as with bi-directional control or symbiont control. In particular, the impact of stochasticity in symbiont number at division is relatively small for holobionts with high symbiont number, such that the correlation only decreases very slightly when symmetric distribution is enforced. Still, when we focus on poor nutrient conditions where holobionts have few symbionts such that symmetric distribution has a large impact, we retrieve the sizer signature that was also found for bi-directional control and symbiont control: without stochasticity at cell division, the correlation between symbiont numbers in consecutive cell cycles is decreased (Fig. S10).

#### Supplementary Tables and Figures

Table S1: Important characteristics of pre-evolved free-living prokaryotes, i.e. cell-cycle efficiency (see Fig. 6) and generalist capacity (plasticity measured as the log-difference between cell-cycle duration at $n = 0.1$ and $n = 100$).

| Strain | Efficiency ($e$) | Plasticity ($\sigma$) |
|---|---|---|
| R8 | 0.476 | 0.965 |
| R9 | 0.402 | 0.866 |
| R2 | 0.307 | -0.355 |
| R3 | 0.160 | 0.497 |

Table S2: Host–symbiont pairs used to initialise evolution experiments (see Table S1 for characteristics of prokaryotes). C1–4 start with identical host and symbiont; C5–6 have different host and symbiont but with very similar phenotypic behavior, i.e. R8 and R9 are both efficient generalists; C7–12 are asymmetric.

| Holobiont | Host | Symbiont |
|---|---|---|
| C1 | R8 | R8 |
| C2 | R9 | R9 |
| C3 | R2 | R2 |
| C4 | R3 | R3 |
| C5 | R8 | R9 |
| C6 | R9 | R8 |
| C7 | R8 | R2 |
| C8 | R2 | R8 |
| C9 | R3 | R2 |
| C10 | R2 | R3 |
| C11 | R8 | R3 |
| C12 | R3 | R8 |

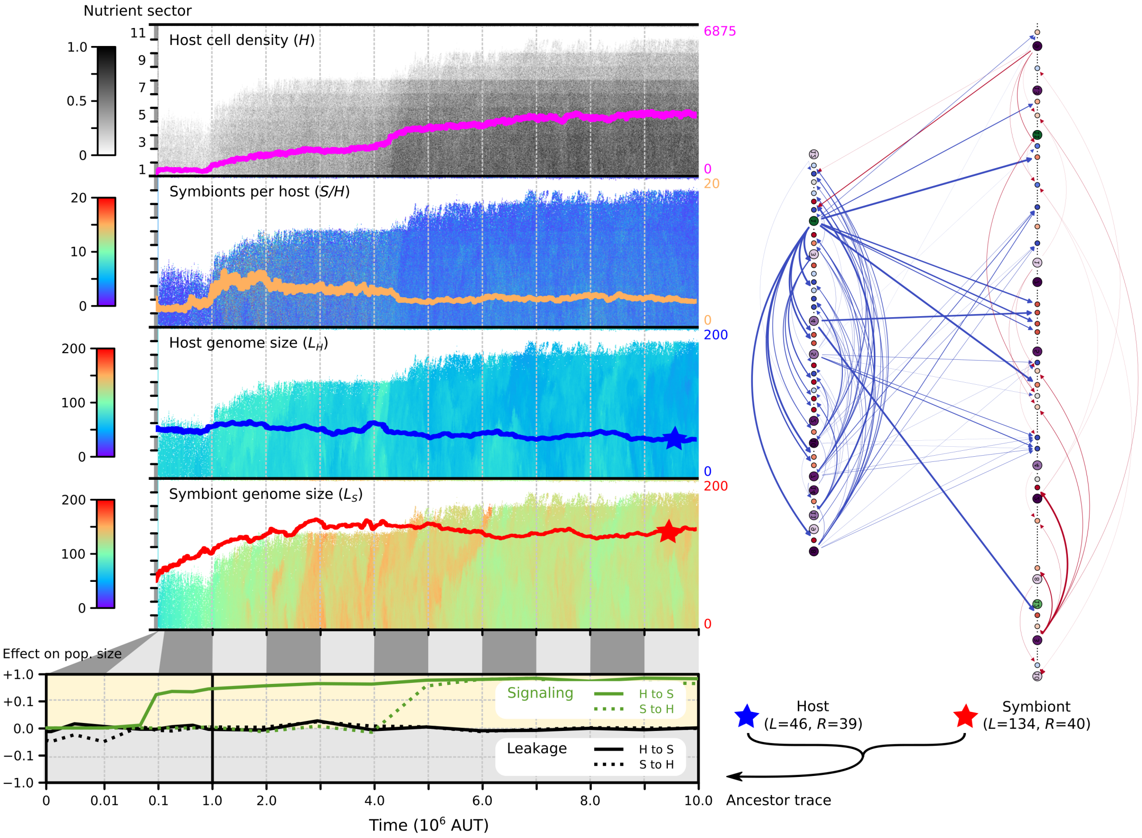


Figure S3: Evolutionary dynamics in a replicate where holobionts evolved bi-directional control (Q3; cf. Fig. 2 in main text). Here, the symbiont evolves a substantially larger genome than the host. Yet, the sizes of their regulatory repertoires are very similar and the host controls more symbiont genes than vice versa. As in Q4, holobionts quickly evolve to be insensitive to leakage (bottom panel) and host-to-symbiont signaling also evolves rapidly. Only from $t = 4 \cdot 10^6$, symbiont-to-host signaling evolves yielding bi-directional control.

### References

von der Dunk, S. H., Hogeweg, P., and Snel, B. (2022). Obligate endosymbiosis explains genome expansion during eukaryogenesis. *bioRxiv*.

Von der Dunk, S. H., Snel, B., and Hogeweg, P. (2022). Evolution of complex regulation for cell-cycle control. *Genome biology and evolution*, 14(5):evac056.

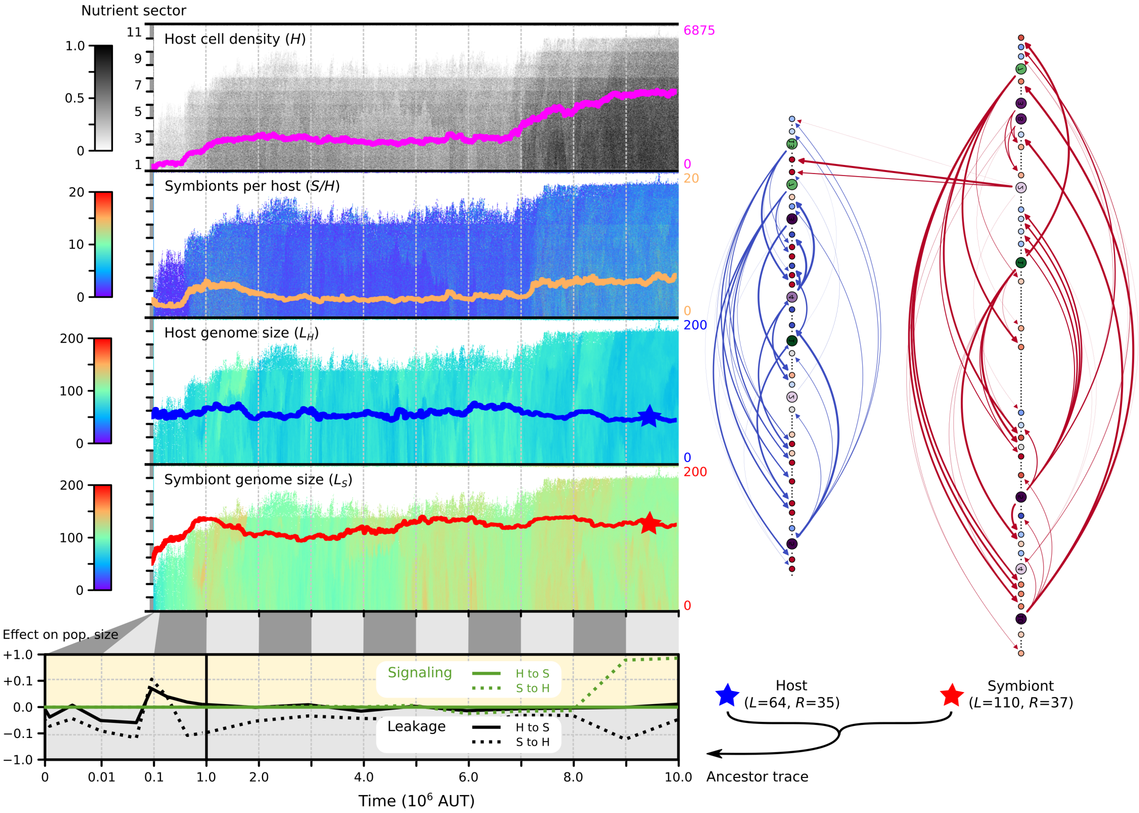


Figure S4: Evolutionary dynamics in a replicate where holobionts evolved symbiont control ($Q_{HS}10$; cf. Fig. 2 in main text). As in Q3, the symbiont evolves a larger genome than the host and the sizes of their regulatory repertoires are comparable. Symbiont-to-host signaling evolves remarkably late, coinciding with substantial population expansion. As before, holobionts quickly evolve to become insensitive to host-to-symbiont leakage (bottom panel). In contrast, holobionts were not exposed to symbiont-to-host leakage, so they remain sensitive to the introduction of this type of leakage.

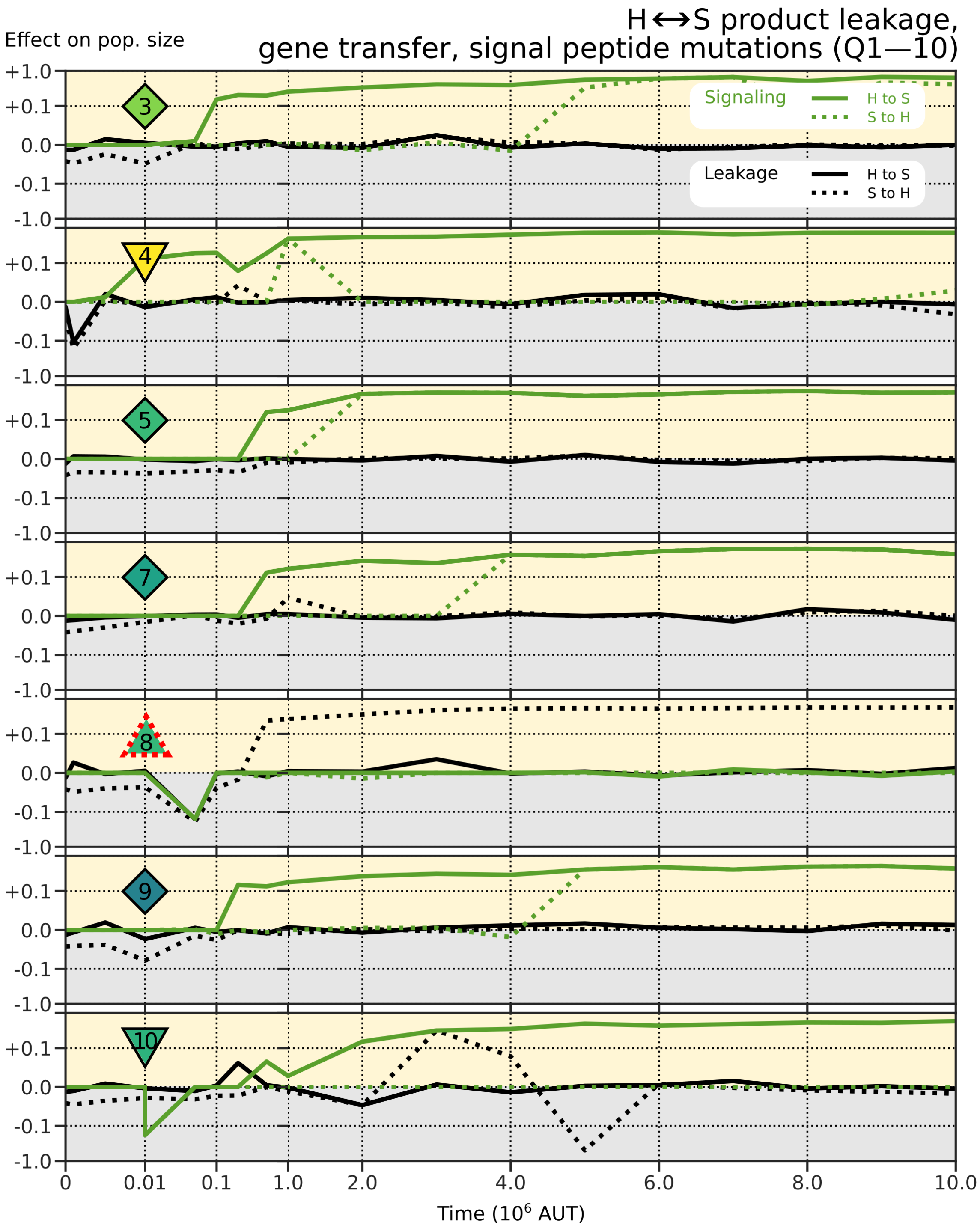


Figure S5: Impact of leakage and signaling along ancestral lineages during evolution with product leakage, gene transfer and signal peptide mutations (Q1–10). Host-to-symbiont signaling always appears early in evolution or does not appear at all. In bi-directional control (Q3, Q5, Q7, and Q9), symbiont-to-host signaling evolves substantially later. In all replicates except Q8, holobionts rapidly become insensitive to leakage. Symbols of replicates correspond to those in Fig. 3 of the main text. Replicates were continued beyond $t = 10 \cdot 10^6$, so we could determine the ancestral lineage up to $t = 10 \cdot 10^6$.

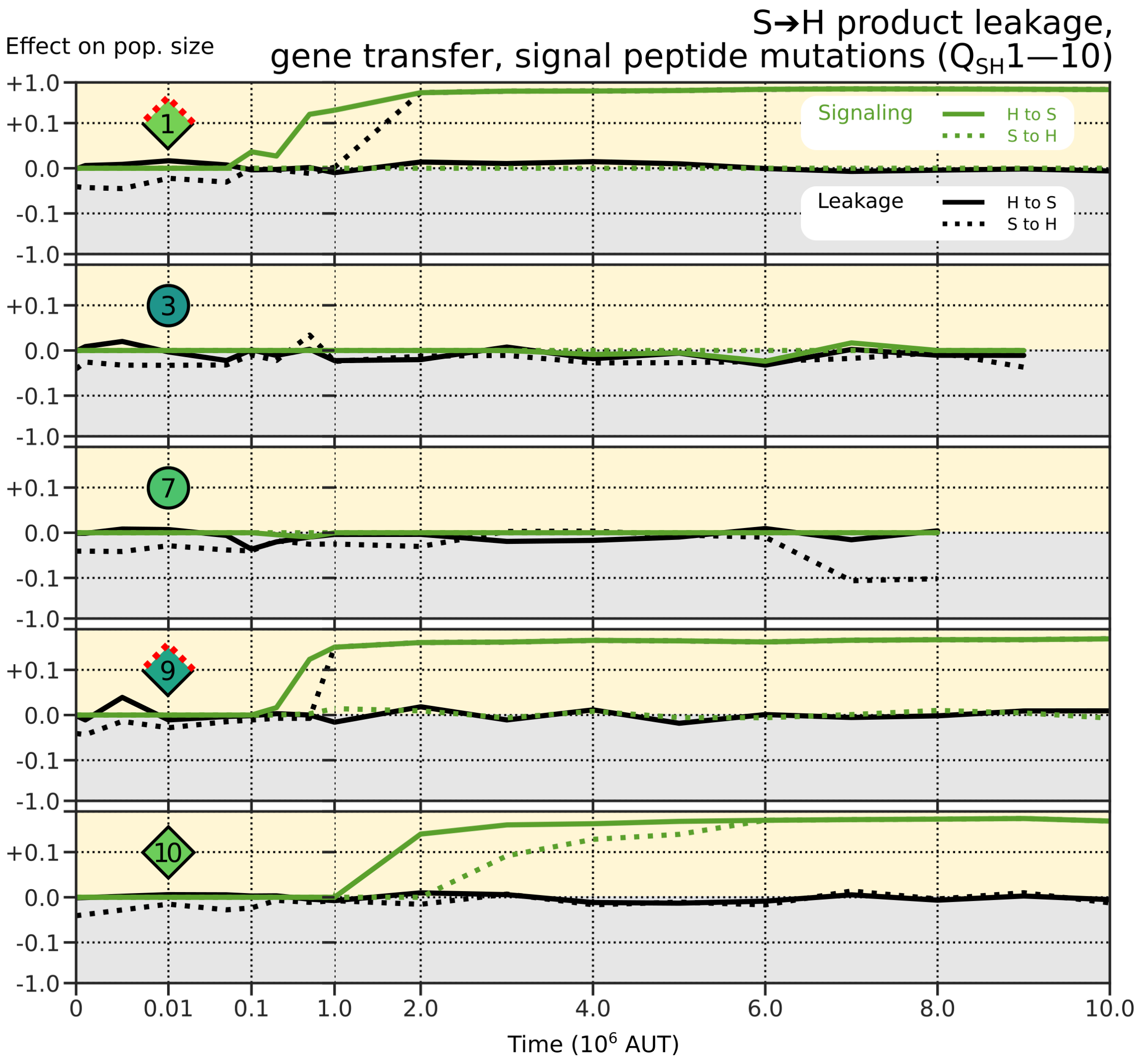


Figure S6: Impact of leakage and targeting along ancestral lineages during evolution with symbiont-to-host product leakage, gene transfer and signal peptide mutations ($Q_{SH}$1–10). As in Fig. S5, host-to-symbiont communication arises before symbiont-to-host communication. Two replicates (Q3 and Q7) do not evolve any functional communication. Symbols of replicates correspond to those in Fig. 3 of the main text. Holobionts were not exposed to host-to-symbiont leakage, and leakage in this direction has little effect on evolved holobionts. Several replicates were continued beyond $t = 10 \cdot 10^6$, so we could determine the ancestral lineage up to $t = 10 \cdot 10^6$.

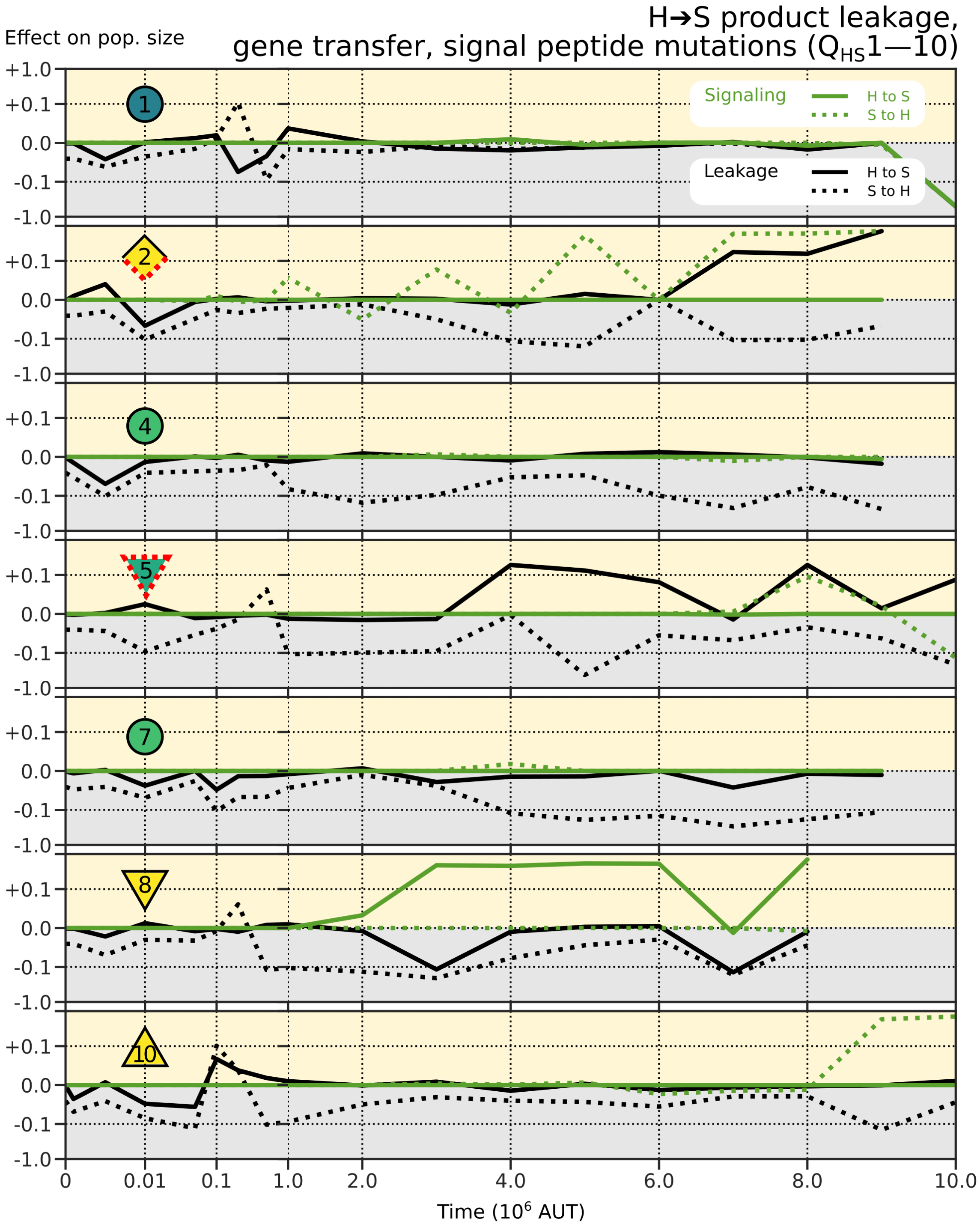


Figure S7: Impact of leakage and targeting along ancestral lineages during evolution with host-to-symbiont product leakage, gene transfer and signal peptide mutations ($Q_{HS}$1–10). Unlike in replicates Q1–10 (Fig. S5), holobionts do not evolve under symbiont-to-host leakage and thus do not become insensitive to it. Symbols of replicates correspond to those in Fig. 3 of the main text. Several replicates were continued beyond $t = 10 \cdot 10^6$, so we could determine the ancestral lineage up to $t = 10 \cdot 10^6$.

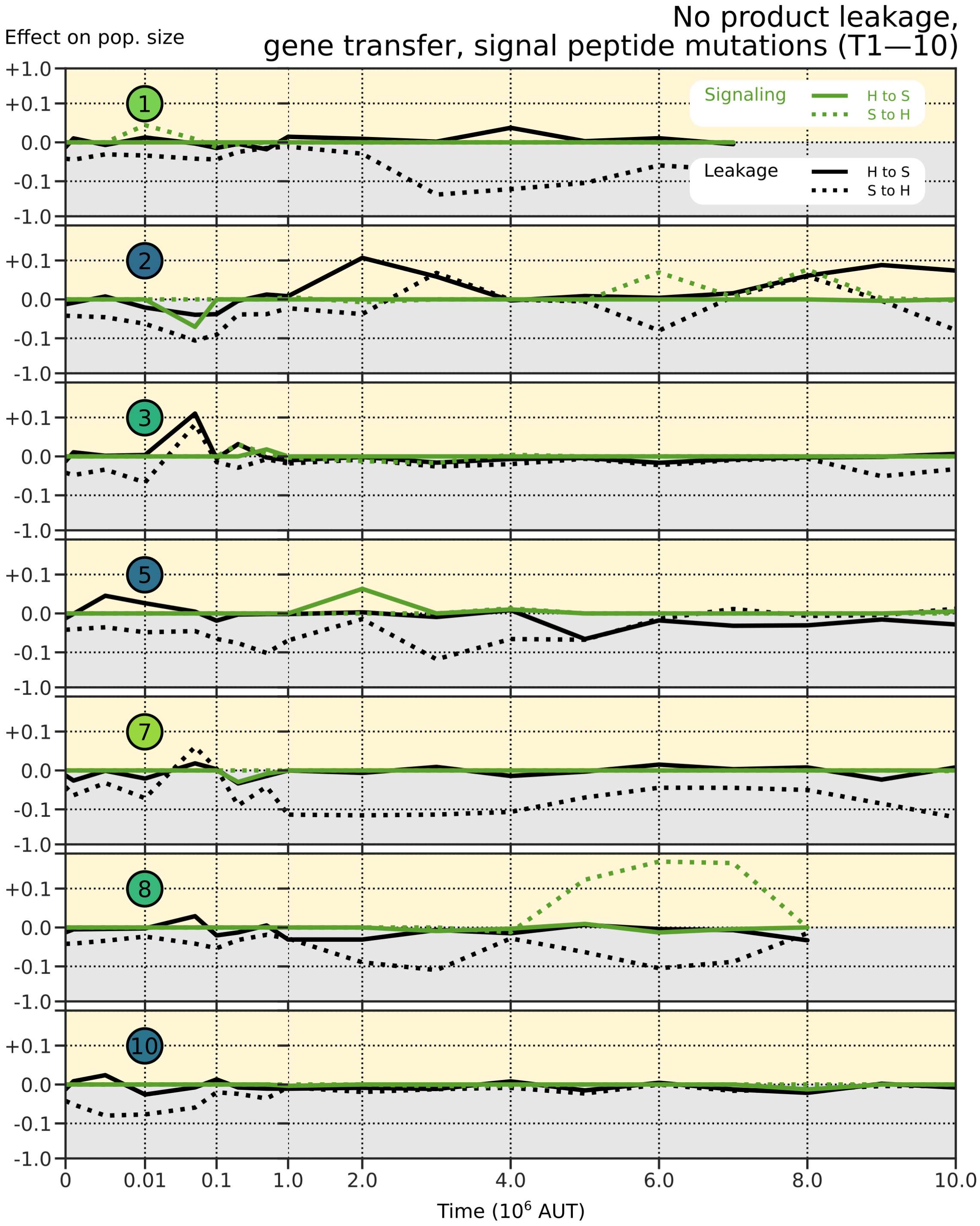


Figure S8: Impact of leakage and targeting along ancestral lineages during evolution with gene transfer and signal peptide mutations but without leakage (T1–10). Functional signaling is only observed transiently in T8. In addition, holobionts do not evolve under leakage, and thus do not become insensitive to it. Still, holobionts might turn out to evolve a transient positive effect of leakage on growth (e.g. early on in T3). Symbols of replicates correspond to those in Fig. 3 of the main text. Several replicates were continued beyond $t = 10 \cdot 10^6$, so we could determine the ancestral lineage up to $t = 10 \cdot 10^6$.

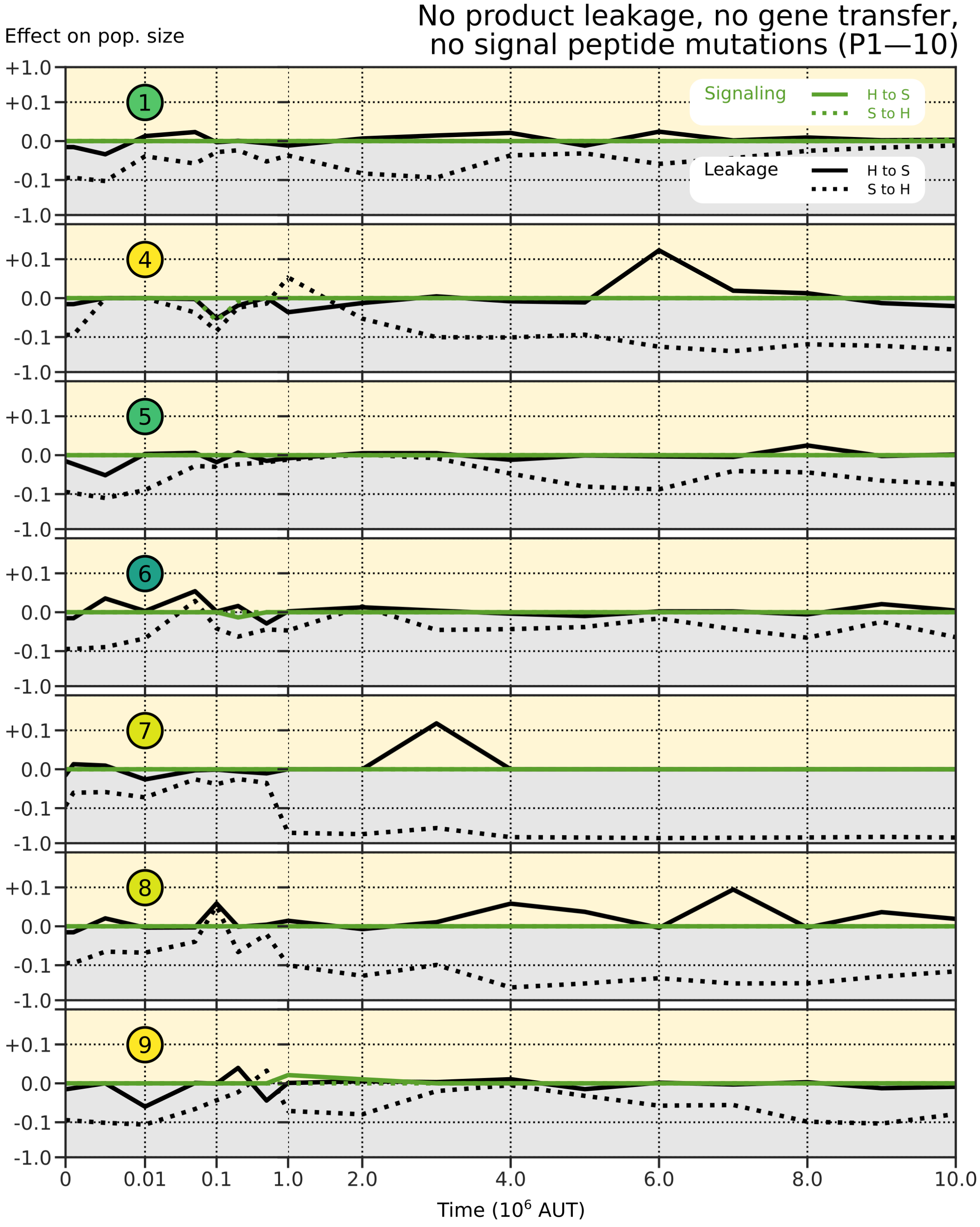


Figure S9: Impact of leakage and targeting along ancestral lineages during evolution without any interference and without the possibility for communication between host and symbiont (P1–10). Symbols of replicates correspond to those in Fig. 3 of the main text. Holobionts were not exposed to leakage, and in many cases remain particularly sensitive to symbiont-to-host leakage. P7 evolves to be especially sensitive to symbiont-to-host leakage, fitting with the large population reduction observed in Fig. 5 in the main text. Conversely, host-to-symbiont leakage is relatively harmless, and even transiently beneficial in P4, P7 and P8. Several replicates were continued beyond $t = 10 \cdot 10^6$, so we could determine the ancestral lineage up to $t = 10 \cdot 10^6$.

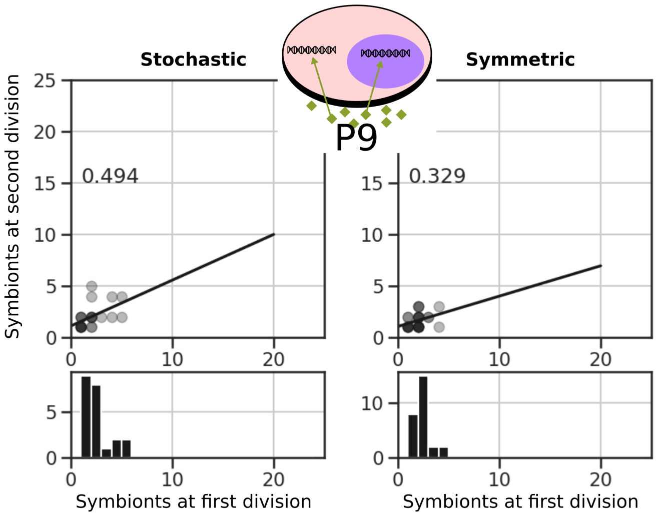


Figure S10: Phenomenological control strategy of P9 becomes more apparent at poor nutrient conditions ($n_{influx} = 5$) where due to low symbiont numbers, the impact of stochasticity at cell division is more prominent (cf. Fig. S2).
